# Arginase 1 is a key driver of immune suppression in pancreatic cancer

**DOI:** 10.1101/2022.06.21.497084

**Authors:** Rosa E. Menjivar, Zeribe C. Nwosu, Wenting Du, Katelyn L. Donahue, Carlos Espinoza, Kristee Brown, Ashley Velez-Delgado, Wei Yan, Fatima Lima, Allison Bischoff, Padma Kadiyala, Daniel Salas-Escabillas, Howard Crawford, Filip Bednar, Eileen Carpenter, Yaqing Zhang, Christopher J. Halbrook, Costas A. Lyssiotis, Marina Pasca di Magliano

## Abstract

An extensive fibroinflammatory stroma rich in macrophages is a hallmark of pancreatic cancer. In this disease, it is well appreciated that macrophages are immunosuppressive and contribute to the poor response to immunotherapy; however, the mechanisms of immune suppression are complex and not fully understood. Immunosuppressive macrophages are classically defined by expression of the enzyme Arginase 1 (Arg1), which we demonstrated is potently expressed in pancreatic tumor associated macrophages from both human patients and mouse models. While routinely used as a polarization marker, Arg1 also catabolizes arginine, an amino acid required for T cell activation and proliferation. To investigate this metabolic function, we used a genetic and a pharmacologic approach to target *Arg1* in pancreatic cancer. Genetic inactivation of *Arg1* in macrophages, using a dual recombinase genetically engineered mouse model of pancreatic cancer, delayed formation of invasive disease, while increasing CD8^+^ T cell infiltration. Treatment of established tumors with the arginase inhibitor CB-1158 exhibited further increased CD8^+^ T cell infiltration, beyond that seen with the macrophage-specific knockout, and sensitized the tumors to anti-PD1 immune checkpoint blockade. Thus, our data demonstrate that Arg1 is more than simply a marker of macrophage function. Rather, Arg1 is also a driver of immune suppression and represents a promising immunotherapeutic target for pancreatic cancer.

## Introduction

Pancreatic ductal adenocarcinoma (PDA) is currently the third leading cause of cancer related deaths in the United States with a five-year survival rate of 11% (Siegel, Miller, Fuchs, & Jemal, 2022). This poor survival rate is due to late detection and ineffective treatments. The hallmark mutation in PDA is in the *KRAS* oncogene, most commonly *KRAS^G12D^* (Hezel, Kimmelman, Stanger, Bardeesy, & Depinho, 2006; Hingorani et al., 2003; Hingorani et al., 2005; Schneider & Schmid, 2003), where disease progression is mediated by loss of tumor suppressor genes, including TP53, SMAD4, and INK4A (Hezel et al., 2006; Maitra & Hruban, 2008).

At current, specific inhibitors for *KRAS^G12D^* are still not clinically available (Liu, Wang, & Li, 2019). The recent clinical implementation of inhibitors for the KRAS^G12C^ mutation in non-small cell lung cancer provide exciting promise for this approach (Canon et al., 2019; Prior, Lewis, & Mattos, 2012; Wilhelm et al., 2006). However, the G12C mutation is exceedingly rare in PDA (Ying et al., 2016). Standard of care for PDA is a combination of systemic chemotherapy that includes FOLFIRINOX or gemcitabine plus nab-paclitaxel (Mizrahi, Surana, Valle, & Shroff, 2020), which only provides limited improvement in patient survival. Although immune checkpoint inhibitors are effective in other cancers, unfortunately this benefit has not translated to PDA (Brahmer et al., 2012; Royal et al., 2010), due to the severely immunosuppressive tumor microenvironment (TME) that characterizes this disease (Morrison, Byrne, & Vonderheide, 2018; Vonderheide & Bayne, 2013).

The pancreatic TME includes cancer associated fibroblasts and a heterogenous population of immune cells, the majority of which are myeloid cells (Gabrilovich & Nagaraj, 2009). These myeloid cells include tumor associated macrophages (TAMs), immature myeloid cells, also referred to as myeloid derived suppressor cells (MDSCs), and granulocytes, such as neutrophils (Gabrilovich, Ostrand-Rosenberg, & Bronte, 2012). In contrast, CD8^+^ T cells are rare in the pancreatic TME, although their prevalence is heterogeneous in different patients (Stromnes, Hulbert, Pierce, Greenberg, & Hingorani, 2017). Single cell sequencing analysis revealed that most CD8^+^ T cells infiltrating PDA have an exhausted phenotype (N. G. Steele et al., 2020). An understanding of the mechanisms mediating immune suppression in pancreatic cancer is needed to design new therapeutic approaches for this disease. Of note, analysis of long-term survivors revealed persistence of tumor-specific memory CD8^+^ T cells, indicating that when an anti-tumor immune response does occur, it leads to effective tumor control (Balachandran et al., 2017); conversely, evidence of loss of antigens over time due to immunoediting has also been described (Luksza et al., 2022). Furthermore, depletion of myeloid cells in mouse models of pancreatic cancer led to activation of anti-tumor T cell responses (Mitchem et al., 2013; Zhang, Velez-Delgado, et al., 2017), spurring an effort to target myeloid cells in pancreatic cancer (Nywening et al., 2016); yet, clinical efficacy has been low, and more approaches are needed.

Myeloid cells infiltrating the neoplastic pancreas, and ultimately those in PDA, express high levels of Arginase 1 (*Arg1*). Indeed, we previously illustrated that *Arg1* expression in myeloid cells is driven by oncogenic *Kras* expression/signaling in epithelial cells, starting during early stages of carcinogenesis (Velez-Delgado et al., 2022; Zhang, Yan, et al., 2017). Beyond our work, Arg1 is widely appreciated as a marker of alternatively polarized macrophages. An increase in tumor myeloid Arg1 expression has been reported in other cancers, including renal cell carcinoma (Rodriguez et al., 2009), breast (Polat, Taysi, Polat, Boyuk, & Bakan, 2003; Singh, Pervin, Karimi, Cederbaum, & Chaudhuri, 2000), colon (Arlauckas et al., 2018), and lung cancer (Miret et al., 2019).

In addition to its role as a marker of alternatively polarized macrophages, ARG1 is also a metabolic enzyme that breaks down the amino acid L-arginine to urea and ornithine (Jenkinson, Grody, & Cederbaum, 1996). Connecting this activity to myeloid Arg1 expression, several reports detail how arginine is necessary for the activation and proliferation of CD8 T cells (Rodriguez, Quiceno, & Ochoa, 2007; Rodriguez et al., 2004). This indicates that myeloid cells may deplete arginine in the TME to damp anti-tumor T cell activity. Based on these concepts, a small-molecule Arginase inhibitor, CB-1158 (INCB001158) (Calithera Biosciences, Inc., South San Francisco, CA), was developed. CB-1158 treatment as monotherapy or in combination with anti-PD-1 checkpoint inhibitor, decreased tumor growth *in vivo* in mice with Lewis lung carcinoma (Steggerda et al., 2017). CB-1158 also inhibits human Arginase and it is being tested in a phase I clinical trial in patients with advanced or metastatic solid tumors (Pham, Liagre, Girard-Thernier, & Demougeot, 2018; Steggerda et al., 2017).

Encouraged by these studies and the prominent disease-specific expression of Arg1 in PDA macrophages, we set forth to determine its functional role. Here, we used a dual recombinase approach to delete *Arg1* in myeloid cell lineages via Cre-loxP technology and at the same time induced oncogenic Kras in pancreatic epithelial cells using the orthogonal Flp-Frt recombination approach. We discovered that deletion of *Arg1* in myeloid cells profoundly reshaped the tumor microenvironment, increased the infiltration of CD8^+^ T cells, and reduced malignant disease progression. However, we also noticed a resistance mechanism whereby other cells in the TME, such as epithelial cells, upregulate Arg1 expression, potentially blunting the effect of its inactivation in myeloid cells. We thus employed a systemic approach, whereby we treated a syngeneic orthotopic model of pancreatic cancer with the Arginase inhibitor CB-1158 and found it to sensitize PDA to anti-PD1 immune checkpoint blockade. These results illustrate for the first time a functional role of Arg1 in pancreatic tumor-derived myeloid cells, reveal novel aspects of intratumoral compensatory metabolism, and provide new inroads to increase the efficacy of checkpoint immune therapy for PDA.

## Results

### Pancreatic cancer infiltrating myeloid cells express Arginase 1

We have previously reported that expression of oncogenic *Kras* in pancreas epithelial cells drives expression of Arginase 1 in macrophages *in vivo* during early stages of carcinogenesis (Velez-Delgado et al., 2022; Zhang, Yan, et al., 2017). We sought to determine whether ARG1 was also associated with late stages of carcinogenesis in mouse and human tumors. As there was no validated antibody for human ARG1 available, we performed *ARG1* RNA in situ hybridization in human PDA, together with co-immunofluorescence staining for the immune cell marker CD45 and for the epithelial cell marker E-cadherin (ECAD) (Figure 1A). We observed prevalent *ARG1* expression in CD45^+^ cells, and occasional low expression in ECAD^+^ cells (Figure 1A). Analysis of our single-cell RNA sequencing (sc-RNA-seq) dataset, which includes 16 human PDA samples (N. G. Steele et al., 2020) (Figure 1B and Figure 1- figure supplement 1A) revealed highest expression of *ARG1* in myeloid cells (Figure 1C). In contrast, we observed minimal expression in CD4^+^ T cells, CD8^+^ T cells, epithelial cells, and fibroblasts (Figure 1C). Thus, we conclude that myeloid cells are the main source of *ARG1* in the pancreatic cancer microenvironment.

**Figure 1.**
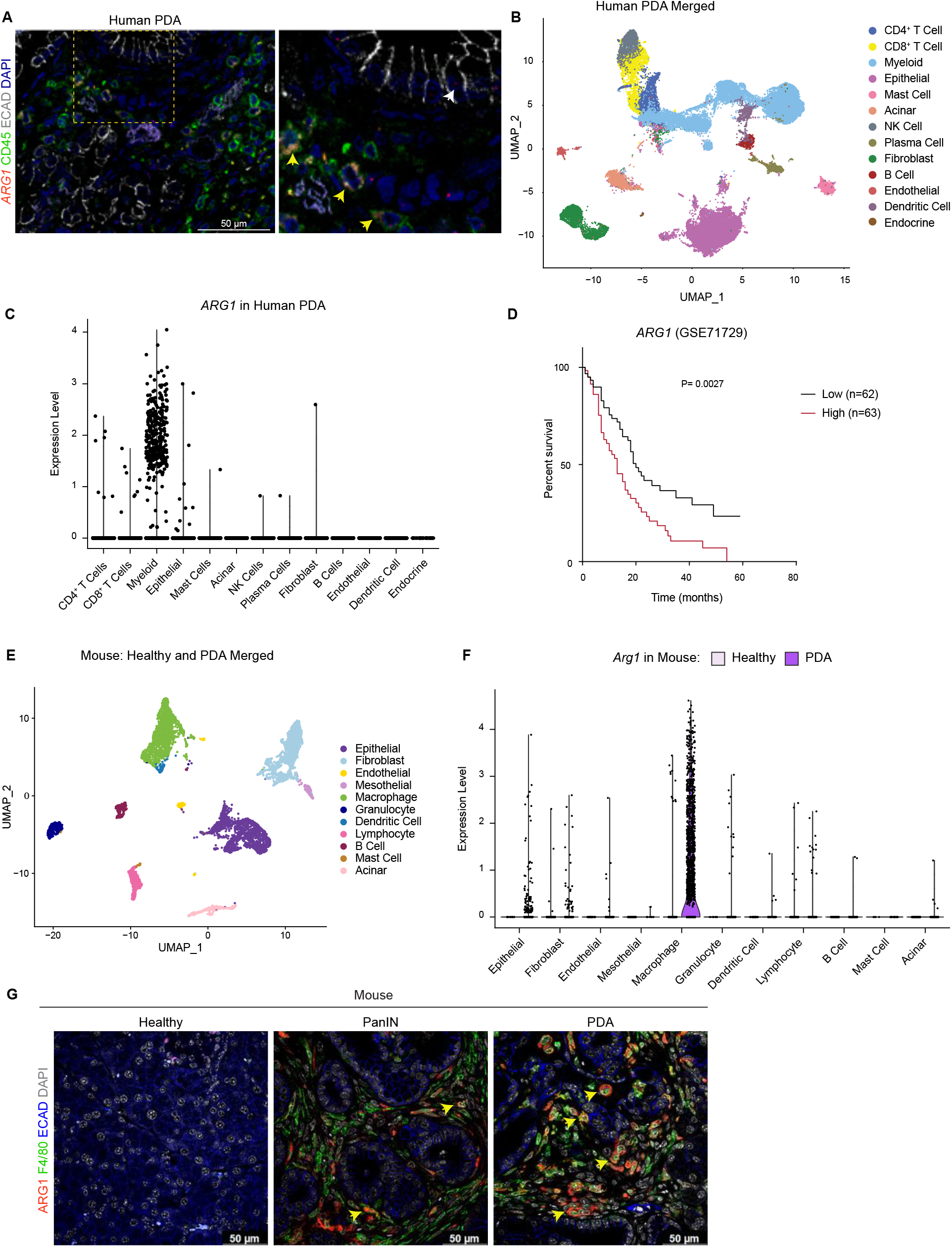
Arginase 1 is highly expressed in human and mouse myeloid cells. (**A**) Representative image of RNA *in situ* hybridization (ISH) of *ARG1* (red) and co-immunofluorescence staining of immune (CD45, green) and epithelial (ECAD, gray) cells in human PDA. Counterstain, DAPI (blue). White arrow points to *ARG1*-ISH in ECAD^+^ cells and yellow arrows point to *ARG1*-ISH in CD45^+^ cells. Scale bar, 50 μm. (**B**) Uniform Manifold Approximation and Projection (UMAP) visualization of 13 identified cell populations from single-cell RNA sequencing (sc-RNA-seq) of 16 human PDA tumors. Data from Steele et al, 2020(N. G. Steele et al., 2020). (**C**) Violin plot of normalized gene expression of *ARG1* in the identified cell populations from the human PDA sc-RNA-seq. (**D**) Survival analysis of a human PDA microarray data set (GSE71729) with low (n=62) and high (n=63) *ARG1* expression. Statistical significance was determined using the Kaplan Meier overall survival Logrank test. (**E**) UMAP visualization of 11 identified populations from healthy and PDA merged mouse sc-RNA-seq. (**F**) Violin plot of normalized gene expression of *Arg1* in the identified cell populations from mouse sc-RNA-seq. (**G**) Representative co-immunofluorescence staining for ARG1 (red), macrophages (F4/80, green), and epithelial (ECAD, blue) cells in mouse tissue at different stages of disease. Counterstain, DAPI (gray). Scale bar, 50 μm. Yellow arrows indicate ARG1 expression in F4/80^+^ cells.

Infiltration of myeloid cells, specifically macrophages, in pancreatic cancer portends worse patient survival (Sanford et al., 2013; Tsujikawa et al., 2017). We thus assessed whether *ARG1* expression correlated with worse patient outcomes. Based on a publicly available human PDA microarray data set (GSE71729) (Moffitt et al., 2015), we found that high *ARG1* expression in pancreatic cancer correlated with worse survival (Figure 1D), suggesting that *ARG1* in myeloid cells may play a functional role in human PDA.

Next, to determine whether the increase in *Arg1* expression was recapitulated in a mouse model of pancreatic cancer, we analyzed our mouse sc-RNA-seq data from healthy mice and tumor-bearing KPC (*LSL-Kras^G12D/+^;LSL-Trp53^R172H/+^;Ptf1a-Cre*) (Hingorani et al., 2005) mice (Figure 1E and Figure 1- figure supplement 1B and 1C). We again detected the highest level of *Arg1* expression in macrophages (Figure 1F and Figure 1- figure supplement 1D). Importantly, the expression was enriched in macrophages from the PDA group compared to macrophages in the normal pancreas (Figure 1F and Figure 1- figure supplement 1D). Other cell types, such as epithelial cells and fibroblasts, only had sporadic *Arg1* expression. To evaluate the protein expression of ARG1, we performed co-immunofluorescence staining. We included healthy pancreas samples, PanIN-bearing pancreata from KC mice (*Ptf1a-Cre;LSL-Kras^G12D/+^*) (Hingorani et al., 2003), and PDA. We did not detect ARG1 protein in the healthy pancreas. In contrast, both PanIN and PDA presented with frequent co-localization of the macrophage marker F4/80 with ARG1, consistent with prevalent expression in this cell population (Figure 1G and Figure 1- figure supplement 1E). Consistent with human data, expression of ARG1 in other cell types was rare (Figure 1G and Figure 1- figure supplement 1E). Thus, both in mouse and human pancreatic cancer ARG1 is highly expressed and largely confined to tumors, where it is predominantly expressed in macrophages.

### Arginase 1 deletion in myeloid cells reduces tumor progression and induces macrophage repolarization

To examine the function of myeloid *Arg1* in PDA, we generated mice lacking *Arg1* expression in myeloid cells. Specifically, we crossed Arg1^f/f^ mice with LysM^Cre/+^ mice to generate LysM^Cre/+^;Arg1^f/f^ mice. LysM^Cre/+^ were generated by inserting the Cre cDNA in the endogenous M lysozyme (LysM) locus (Clausen, Burkhardt, Reith, Renkawitz, & Forster, 1999), which is broadly expressed in myeloid cells, including macrophages and neutrophils (Clausen et al., 1999; El Kasmi et al., 2008). To validate the deletion of *Arg1* in macrophages, we harvested bone marrow cells from wild type (WT) or LysM^Cre/+^;Arg1^f/f^ mice and cultured these directly in pancreatic cancer cell conditioned media (CM) for 6 days. This protocol yields bone marrow-derived TAMs (Figure 2A), as previously described (Zhang, Velez-Delgado, et al., 2017). ARG1 expression was readily detectable in wild type TAMs; in contrast, LysM^Cre/+^;Arg1^f/f^ TAMs had none or very low expression, indicating efficient Cre recombination (Figure 2B). We then sought to determine whether ARG1 expression in TAMs affected the metabolite composition of the media. We thus performed extracellular metabolomics by liquid chromatography couple tandem mass spectrometry (LC-MS/MS) and observed elevated levels of L-Arginine in the LysM^Cre/+^;Arg1^f/f^ TAM medium compared with WT TAM medium (Figure 2C). This finding is consistent with wild type TAMs depleting Arginine from their growth medium at an enhanced rate, relative to TAMs lacking *Arg1* expression.

**Figure 2.**
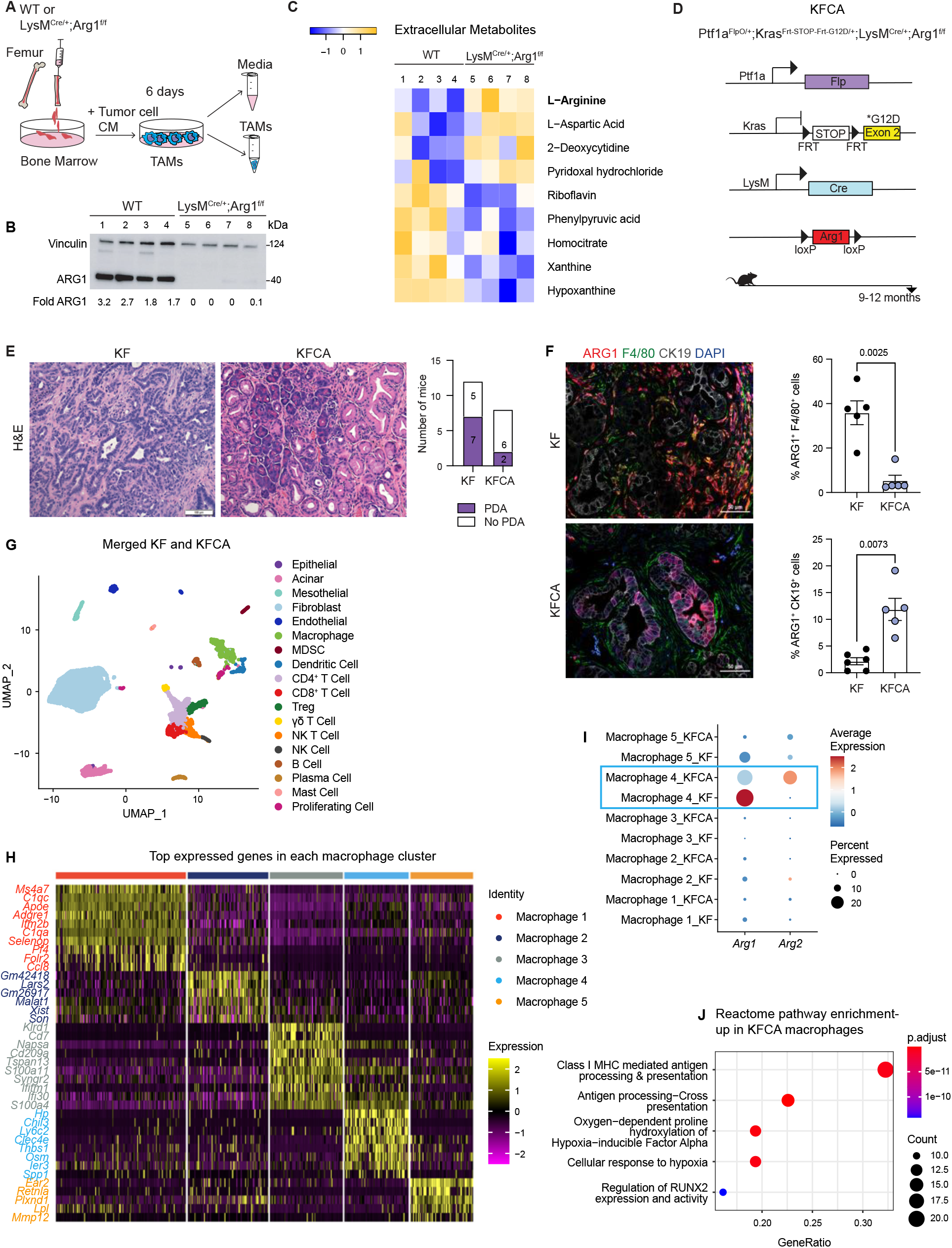
Arginase 1 deletion in myeloid cells reduces tumor formation and induces macrophage repolarization in a spontaneous PDA mouse model. (**A**) Schematic illustration for the generation of tumor associated macrophages (TAMs) from WT and LysM^Cre/+^;Arg1^f/f^ mice. (**B**) Representative image of western blot for ARG1 levels in WT and LysM^Cre/+^;Arg1^f/f^ TAMs. Vinculin, loading control. (**C**) Heatmap of statistically significantly different extracellular metabolites from WT (lanes 1-4) and LysM^Cre/+^;Arg1^f/f^ (lanes 5-8) TAM media. (**D**) Genetic makeup of the Ptf1a^FlpO/+^;Kras^Frt-STOP-Frt-G12D/+^; LysM^Cre/+^;Arg1^f/f^ (KFCA) mouse model for the deletion of *Arg1* in myeloid cells during PDA tumorigenesis. Data shown here from mice aged to 9-12 months, n= 8-12/group. (**E**) Representative H&E staining from age matching KF and KFCA mice. Scale bar, 100 μm. Histopathology evaluation shown on the right. (**F**) Representative image of co-immunofluorescence staining for ARG1 (red), macrophages (F4/80, green), epithelial (CK19, gray), and DAPI (blue) in KF and KFCA tissue. Scale bar, 50 μm. Quantification on the right, n= 5-6/group (**G**) UMAP visualization for the identified cell populations in merged KF and KFCA sc-RNA-seq. (**H**) Heatmap of top differentially expressed genes in the macrophage subclusters identified from KF and KFCA pancreata. (**I**) Dot plot visualization of *Arg1* and *Arg2* expression in KF and KFCA macrophage clusters. Average expression is shown by color intensity and expression frequency by dot size. (**J**) Reactome pathway enrichment analysis showing significantly upregulated pathways in KFCA macrophages.

To investigate the function of *Arg1* in myeloid cells during PDA progression, we generated Ptf1a^FlpO/+^;Kras^Frt-STOP-Frt-G12D/+^ (KF); LysM^Cre/+^;Arg1^f/f^ mice, hereafter referred to as KFCA (Figure 2D and Figure 2- figure supplement 1A). The dual-recombinase system integrates both *Flippase-FRT* (*Flp-FRT)* and *Cre-loxP* recombination technologies to independently modify epithelial cells and myeloid cells (Garcia et al., 2020; Wen et al., 2019). We aged a cohort of KF and KFCA mice for two months, a timepoint at which PanIN lesions are formed in KF mice. To evaluate the efficiency of *Arg1* deletion, we performed co-immunofluorescent staining for ARG1, F4/80, and CK19. As expected, we observed abundant expression of ARG1 in F4/80^+^ macrophages in KF, and minimal to no expression of ARG1 in macrophages from KFCA tissue (Figure 2- figure supplement 1B). We then performed histopathological evaluation of the tissue to determine the functional effect of loss of myeloid *Arg1*. Here, we observed a decrease in ADM and PanIN lesions, accompanied by a reduction of desmoplastic stroma in KFCA, compared to age-matched KF pancreata (Figure 2- figure supplement 1C).

To investigate the causes underlying the reduction in PanIN formation, we performed an in-depth histological analysis. We previously showed that macrophages can directly promote epithelial cell proliferation during early stages in carcinogenesis (Zhang, Yan, et al., 2017). We and others have also demonstrated that macrophages also indirectly promote carcinogenesis by suppressing CD8^+^ T cell infiltration and activation (Bayne et al., 2012; Mitchem et al., 2013; Pylayeva-Gupta, Lee, Hajdu, Miller, & Bar-Sagi, 2012; Zhang, Velez-Delgado, et al., 2017; Zhu et al., 2017). Immunostaining for the proliferation marker Ki67 revealed a reduction in proliferation, a reduction in cell death as determined by Cleaved caspase-3 (CC3) staining, and no changes in infiltrating F4/80^+^ macrophages or CD8^+^ T cells (Figure 2- figure supplement 1D). Thus, loss of Arg1 in macrophages appears to reduce the ability of macrophages to promote proliferation but, at this stage, does not appear to correlate with changes in the immune system. These findings are consistent with our previous observations on the role of macrophages during the onset of carcinogenesis (Zhang, Yan, et al., 2017).

Given the reduction in PanIN formation, we next investigated the effects of myeloid *Arg1* deletion on progression to PDA. For this purpose, we aged KF and KFCA mice to 9-12 months (Figure 2D), an age where we have previously observed invasive cancer formation in KF mice (Garcia et al., 2020). Accordingly, 7 out of 12 KF mice (58%) had invasive disease; in contrast, only 2 out of 8 KFCA mice had progressed (25%) (Figure 2E). We then stained the tissue for ARG1 together with F4/80, and CK19. As expected, we observed high ARG1 expression in macrophages in KF tissues and little to no ARG1 expression in macrophages in KFCA pancreata (Figure 2F). Surprisingly, we observed substantial ARG1 expression in epithelial cells from the KFCA pancreata, while epithelial cells in KF mice had little to no ARG1 expression (Figure 2F and Figure 2- figure supplement 2A). So, while deletion of *Arg1* in myeloid cells impairs malignant progression, it unleashes a compensatory upregulation of *Arg1* in epithelial cells.

We noted that the epithelia that expressed ARG1 in KFCA mice had an elongated appearance, consistent with tuft cells, a cell type that is not present in the healthy pancreas but common in low grade PanINs (Delgiorno et al., 2014). By co-immunostaining for the tuft cell marker COX1, we confirmed that tuft cells are the exclusive source of epithelial ARG1 in these samples (Figure 2- figure supplement 2A), suggesting a tuft cell response to accumulating arginine in the microenvironment.

To comprehensively characterize the phenotype of myeloid *Arg1* deleted mice, we performed sc-RNA-seq on KF and KFCA pancreata dissected from 11 months old mice. Unsupervised clustering identified abundant stromal and immune cells, and a small population of epithelial cells, in both genotypes (Figure 2G and Figure 2- figure supplement 2B and 2C). The percentage of total macrophages was similar between KF and KFCA pancreas (Figure 2- figure supplement 2D, 2E, and 2F). We then subclustered the macrophages, from which we identified five different macrophage populations based on distinct gene profiles (Figure 2H and Figure 2- figure supplement 2G). The macrophage 1 population was defined by *Apoe, C1qa, and C1qc,* markers of tumor associated macrophages (TAMs) in mouse and human pancreatic cancer (Kemp, Carpenter, et al., 2021), while the macrophage 5 population expressed *Ear2* and *Retnla* (Figure 2H). Since macrophage APOE is tumor-promoting (Kemp, Carpenter, et al., 2021), we performed co-immunofluorescent staining for APOE, F4/80, and ECAD and detected a reduction in APOE expression (Figure 2- figure supplement 2H).

Intriguingly, sc-RNA-seq revealed that deletion of *Arg1* in myeloid cells led to an upregulation of *Arg2* expression, mainly in the macrophage 4 population expressing *Chil3*, *Ly6c2*, and *Clec4e* genes (Figure 2I), suggesting the existence of a compensatory mechanism. We then performed reactome pathway enrichment analysis on the macrophage sc-RNA-seq data to determine both up- and down-regulated pathways (Figure 2J and Figure 2- figure supplement 2I). Interestingly, KFCA macrophages exhibited upregulation of signaling pathways involved in MHC I antigen processing and cross presentation (Figure 2J), suggesting an improvement in antigen specific CD8^+^ T cell activation upon *Arg1* deletion.

### Deletion of Arginase 1 in myeloid cells increases CD8^+^ T cell infiltration and activation

Since single cell RNA sequencing suggested increased immune activity and given that macrophages are known to inhibit anti-tumor T cell responses (DeNardo & Ruffell, 2019), we stained KF and KFCA pancreata for CD8 and observed increased CD8^+^ T cell infiltration in the latter (Figure 3A). We then sub-clustered T and NK cells in the sc-RNA-seq datasets from KF and KFCA pancreata and classified these into the following subclusters: naïve CD8^+^ T cells, cytotoxic CD8^+^ T cells, exhausted CD8^+^ T cells, CD4^+^ T cells, regulatory T cells (Treg), γδ T cells, natural killer T (NKT) and natural killer (NK) cells (Figure 3B and 3C). From this analysis, we observed an increase in the percentage of both cytotoxic and exhausted CD8^+^ T cells in the KFCA model compared with KF (Figure 3D and 3E). Correspondingly, expression of genes related to CD8^+^ T cell cytotoxicity, including *Granzyme b* (*Gzmb*), *Perforin 1* (*Prfn1*), and *Interferon gamma* (*Ifng*) were upregulated in CD8^+^ T cells from the KFCA model (Figure 3F). Accordingly, co-immunofluorescent staining for CD8, GZMB, and ECAD showed an increase in GZMB-expressing CD8^+^ T cells in KFCA pancreata (Figure 3G). We also found that the expression of genes involved in T cell exhaustion such as *Cytotoxic T-lymphocyte associated protein 4* (*Ctla4*), *Furin*, *Lymphocyte activating 3* (*Lag3*), and *Programmed cell death 1* (*Pdcd1*) were upregulated in the KFCA model, compared with the KF (Figure 3H). Taken together, deletion of *Arg1* in myeloid cells resulted in an increase in CD8^+^ T cell infiltration and activation, counterbalanced by an increase in exhaustion as well.

**Figure 3.**
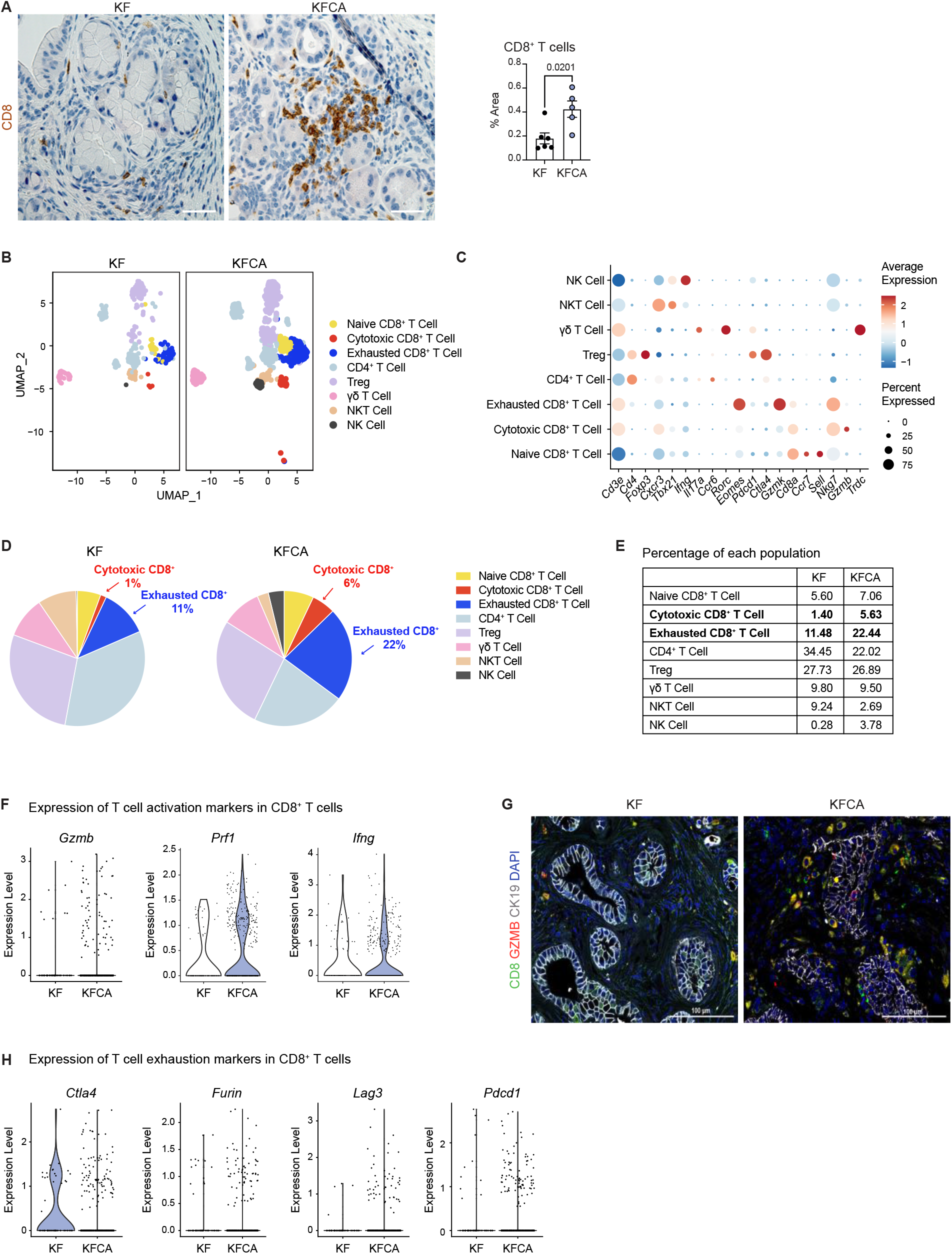
Arginase 1 deletion in myeloid cells increases CD8^+^ T cell infiltration and activation in a spontaneous PDA mouse model. (**A**) Representative images of CD8 immunohistochemistry staining (brown) in KF and KFCA tissue. Scale bar, 50 μm. Quantification of positive area on the right, n=5-6/group. Student’s t test was used to determine statistical significance. (**B**) UMAP visualization of defined T and NK cell clusters comparing sc-RNA-seq data from KF and KFCA. (**C**) Dot plot of lineage markers used to identify the different types of lymphocytes. Dot size shows expression frequency, dot color shows average expression. (**D**) Pie charts showing the proportion of the identified lymphocyte populations in KF and KFCA sc-RNA-seq, percentage values are provided for populations that differ dramatically between KF and KFCA. (**E**) Table showing the percentage of each identified lymphocyte population in KF and KFCA sc-RNA-seq. (**F**) Violin plots showing normalized expression levels of T cell activation markers in all CD8^+^ T cell populations identified in KF and KFCA sc-RNA-seq data. (**G**) Representative images of co-immunofluorescence staining for CD8 (green), GZMB (red), CK19 (gray) and DAPI (blue). Scale bar, 100 μm. (**H**) Violin plots showing normalized expression levels of T cell exhaustion markers in all CD8^+^ T cell populations identified in KF and KFCA.

### Systemic Arginase inhibition in combination with anti-PD1 immune checkpoint reduces tumor growth

Our genetic model revealed that deletion of *Arg1* in myeloid cells reduced PanIN/PDA progression and was accompanied by an increased infiltration of CD8^+^ T cells. However, we also observed several compensatory mechanisms that may have blunted the effect of *Arg1* loss. These included expression of *Arg2* in myeloid cells and expression of *Arg1* in epithelial cells. We reasoned that systemic inhibition of Arginase (Steggerda et al., 2017), using a pharmacologic approach might bypass these compensatory mechanisms.

We implanted a KPC pancreatic cancer cell line (7940B) (Long et al., 2016) orthotopically into the pancreas of syngeneic C57BL6/J mice. Upon tumor detection (by palpation or by ultrasound), we randomly divided the mice into two groups to receive either vehicle or Arginase inhibitor (CB-1158, Calithera Biosciences) (Figure 4A). We pharmacologically treated the mice for 10 days and then harvested the tumors 20 days post-implantation.

**Figure 4.**
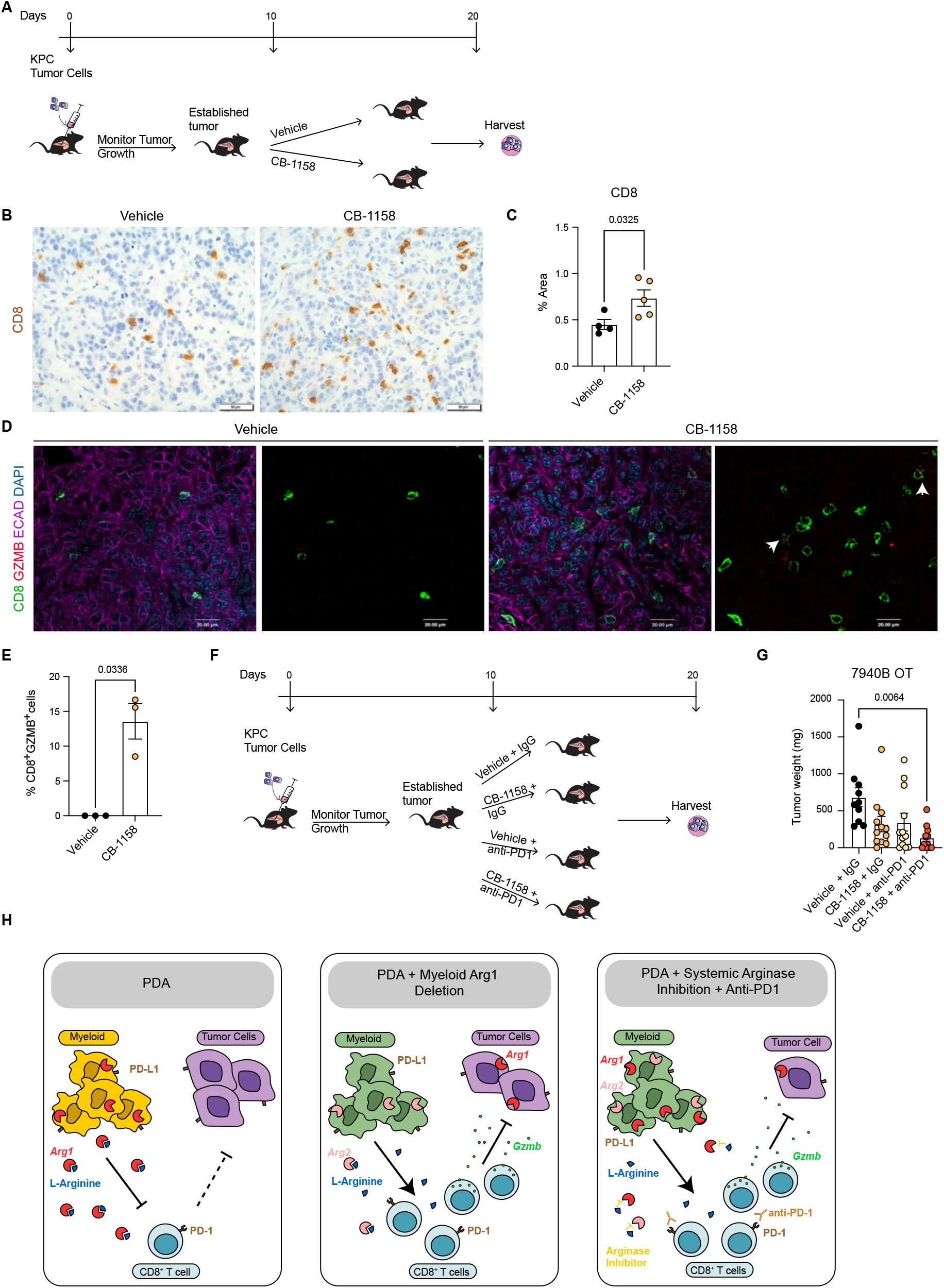
Systemic inhibition of Arginase by CB-1158 in combination with anti-PD1 reduce tumor growth in an orthotopic PDA mouse model. (**A**) Experimental timeline and design for orthotopic transplantation of 7940B KPC cells in syngeneic mice. (**B**) Representative images of immunohistochemistry staining for CD8 (brown). Scale bar, 50 μm. (**C**) Quantification of positive area of CD8 staining from **B.** Statistical significance was determined using unpaired t-test with Welch’s correction. (**D**) Representative images of co-immunofluorescence staining for CD8 (green), GZMB (red), ECAD (purple) and DAPI (blue) in the vehicle and CB-1158 treated group. Scale bar, 20 μm. White arrows point to co-localization of CD8 (green) with GZMB (red). (**E**) Percentage of CD8^+^ and GZMB^+^ cells. Statistical significance was determined using unpaired t-test with Welch’s correction. (**F**) Experimental timeline and design for orthotopic transplantation of 7940B KPC cells in syngeneic mice and different treatment groups. (**G**) Final tumor weight (mg) from the different treatment groups, n= 10-12/group. Statistical significance was determined using two-way ANOVA with Tukey’s multiple comparisons correction test. (**H**) Diagram depicting our working model. First panel, Arg1 in PDA; second panel, deletion of Arg1 in myeloid cells in a spontaneous PDA mouse model; and third panel, PDA with systemic Arginase inhibition in combination with anti-PD1 immune checkpoint blockade.

First, we investigated the infiltration of CD8^+^ T cells by staining the vehicle and CB-1158 mouse tissue by immunohistochemistry for CD8. Interestingly, we observed that systemic inhibition of Arginase by CB-1158 increased the infiltration of CD8^+^ T cells (Figure 4B and 4C), recapitulating our findings from the genetically engineered model. To identify whether there was an increase in CD8^+^ T cell activation, we stained the tissue samples for CD8 and the activation marker GZMB, as well as the epithelial marker ECAD. While in the control tissue GZMB expression was rare, in the CB-1158 treatment group GZMB was common in CD8^+^ T cells (Figure 4D and 4E). Data from the genetic model showed an increase in T cell activation, but also an increase in exhaustion. We thus repeated the syngeneic orthotopic transplantation experiment described above, but this time we added blockade of the PD1 immune checkpoint to circumvent T cell exhaustion. Tumor bearing mice were divided into four different groups to receive: (1) vehicle + IgG control, (2) Arginase inhibitor (CB-1158) + IgG, (3) vehicle + anti-PD1, and (4) CB-1158 + anti-PD1 (Figure 4F). We observed a trending decrease in tumor weight in the CB-1158 + IgG group and a significant decrease in tumor weight in the combination group of CB-1158 + anti-PD1 immune checkpoint blockade compared with our vehicle control group (Fig. 4f). These findings recapitulated the decrease in tumor formation observed in the KFCA mouse model.

We then proceeded to characterize the tumor tissue. We examined ARG1 expression in macrophages (F4/80^+^) and epithelial (ECAD^+^) cells by co-immunofluorescence staining and observed no changes in ARG1 distribution or expression among treatment groups (Figure 4- figure supplement 1A), consistent with the notion that the CB-1158 Arginase inhibitor does not affect enzyme production but suppresses its activity. H&E staining showed large necrotic areas in both CB-1158 and CB-1158 + anti-PD1 groups (Figure 4- figure supplement 1B). To further characterize the tumor tissue, we stained for cell proliferation (Ki67) and apoptotic cell death (CC3) by immunohistochemistry staining (Figure 4- figure supplement 1C and 1D). The combination treatment group showed a significant decrease in cell proliferation compared to the vehicle control group (Figure 4- figure supplement 1C and 1E). We also observed either a significant or trending increase in CC3 upon CB-1158 + IgG, vehicle + anti-PD1, or the combination treatment (Figure 4- figure supplement 1D and 1F). Together, these findings indicate that Arginase inhibition in combination with anti-PD1 reduces immune suppression and decreases pancreatic cancer tumor growth (See working model in Figure 4H).

## Discussion

Abundant myeloid cells in the TME are associated with poor prognosis in multiple types of cancer, including PDA (Gentles et al., 2015) (Sanford et al., 2013) (Tsujikawa et al., 2017). In contrast, high levels of T cells correlate with longer survival (Gentles et al., 2015) (Balachandran et al., 2017). We and others have shown that myeloid cells promote pancreatic cancer growth both directly and by inhibiting CD8^+^ T cell anti-tumor immunity (Zhang, Yan, et al., 2017) (Zhang, Velez-Delgado, et al., 2017) (Mitchem et al., 2013) (Halbrook et al., 2019) (C. W. Steele et al., 2016) (Liou et al., 2015). The specific mechanisms by which macrophage drive immune suppression, and how to best target macrophages to improve outcomes in pancreatic cancer remain poorly understood.

Macrophages are plastic cell types, traditionally classified into pro-inflammatory “M1” and anti-inflammatory M2” subtypes, based on their gene expression pattern (Mantovani, Sozzani, Locati, Allavena, & Sica, 2002). Recently, a wealth of evidence now supports the notion that tumor associated macrophages (TAMs) are distinct from M1 and M2 (Boyer et al., 2022; DeNardo & Ruffell, 2019; Halbrook et al., 2019). Adding further complexity, TAMs also exhibit heterogeneous populations within an individual tumor. ARG1 is classically considered a “marker” of the anti-inflammatory M2 state. In this study, we show that the enzyme Arginase 1 (ARG1) is expressed in human and mouse TAMs, as well as in other myeloid populations. Further, when we stratified human PDA based on *ARG1* expression, we found inverse correlations between *ARG1* and survival. However, prior to this work, a functional role of Arg1 in pancreatic TAMs and tumor immunity had not been evaluated.

Arginases are enzymes that hydrolyze the amino acid L-Arginine to urea and L-ornithine in the liver urea cycle (Jenkinson et al., 1996). An analysis of plasma metabolites and tumor interstitial fluid in an autochthonous PDA mouse model revealed that L-Arginine was drastically reduced in the tumor interstitial fluid (Sullivan et al., 2019). Additionally, ornithine levels increased in the PDA tumor interstitial fluid compared to plasma (Sullivan et al., 2019), suggesting Arginase contribution to immunosuppression through depletion of Arginine.

There are two isoforms of Arginase, ARG1 and ARG2, located in the cytoplasm and mitochondria, respectively (Gotoh et al., 1996; Munder et al., 2005). The *Arginase* genes share a 58% sequence identity (Haraguchi et al., 1987) and are almost identical at the catalytic site. *Arg1* is normally expressed by hepatocytes, but it is also expressed in TAMs in a variety of tumor types. *Arg2* is located in various cell types, including renal cells, neurons, and macrophages (Bronte & Zanovello, 2005; Caldwell, Rodriguez, Toque, Narayanan, & Caldwell, 2018). In a mouse model of Lewis lung carcinoma, myeloid cells express high levels of *Arg1*, resulting in impaired T cell function (Rodriguez et al., 2004). In that study, 3LL tumor cells were implanted subcutaneously in the right flank of the mice and at the same time, mice were treated with the pan-Arginase inhibitor tool compound N-hydroxy-nor-L-Arg. *Arg1* inhibition reduced subcutaneous tumor growth in a dose dependent manner. The anti-tumor effect of the Arginase inhibitor was in part dependent on T cell function, as inhibition on tumor growth was not observed when mice lacking a functional immune system were treated with the Arginase inhibitor (Rodriguez et al., 2004). In a more recent study (Miret et al., 2019), inhibition of Arginase activity by the Arginase inhibitor tool compound Cpd9 (Van Zandt et al., 2013) decreased T cell suppression and reduced tumor growth in a Kras^G12D^ lung cancer mouse model.

To address the role of macrophage ARG1 in PDA, we used a dual recombinase, genetic approach to delete *Arg1* in myeloid cells and aged the mice until most animals in the control group developed invasive PDA. We discovered that deletion of *Arg1* in myeloid cells reduced progression to invasive disease, conversely resulting in accumulation of early lesions with prominent Tuft cells, a cell type that is protective towards tumor progression (DelGiorno et al., 2020; Hoffman et al., 2021). Reduced tumor progression was accompanied by changes in the immune microenvironment, such as an increase in infiltration and activation of CD8^+^ T cells. However, a fraction of the mice developed invasive disease. Complete CD8^+^ T cell reactivation against tumors did not occur as we also observed an increase in exhausted CD8^+^ T cells. Additionally, we observed increased *Arg2* expression in myeloid cells. This finding is in line with a report in which patients with ARG1 deficiency were also found to have an increase in ARG2 levels (Crombez & Cederbaum, 2005; Grody et al., 1993), suggesting compensation of ARG2 for the loss of ARG1.

Further analysis revealed that, in the absence of myeloid *Arg1*, the chemosensory tuft cells began to express ARG1. Tuft cells sense their surrounding environment and respond in a variety of ways, including modifying immune responses in a way that restrains PDA malignant progression (DelGiorno et al., 2020; Hoffman et al., 2021). Their compensation for the loss of myeloid *Arg1* expression in what is likely now an arginine-rich microenvironment is consistent with these functions and suggests that systemic inhibition of ARG1 may have a more profound effect than its ablation from myeloid cells alone.

To systemically inhibit ARG1 we used CB-1158 (INCB001158), an orally bio-available small molecule inhibitor of Arginase. CB-1158 is not cell permeable, and thus does not inhibit liver ARG1, which would lead to immediate toxicity (Steggerda et al., 2017). *In vitro*, CB-1158 inhibits human recombinant ARG1 and ARG2. The *in vivo* action is attributed to inhibition of extracellular ARG1 released by TAMs and other myeloid cells (Steggerda et al., 2017). We used a syngeneic, orthotopic transplantation model based on KPC pancreatic cancer cells transplanted in C57Bl6/J mice (Hingorani et al., 2005; Long et al., 2016). Treatment with CB-1158 led to an increase in infiltrating CD8^+^ T cells, recapitulating the findings in the spontaneous model. Immune checkpoint therapies such as PD-1/PD-L1 used to enhance the anti-tumor immune response of T cells are not effective in pancreatic cancer (Brahmer et al., 2012), in part due to resistance mechanisms induced by myeloid cells. We found that the combination treatment of CB-1158 with anti-PD1 immune checkpoint blockade reactivated exhausted CD8^+^ T cells and decreased tumor growth. CB-1158 is currently evaluated as a single agent and in combination with immune checkpoint therapy or chemotherapy in patients with solid tumors (www.clinicaltrials.gov, NCT02903914 and NCT03314935).

Because of the heterogeneity of myeloid cells acquired by their interaction with the tumor microenvironment, myeloid cells most likely utilize multiple mechanisms that affect their function. Our findings show that one of the mechanisms by which myeloid cells promote immune suppression and tumor growth in pancreatic cancer is through over expression and activity of *Arg1*. Thus, Arginase inhibition may be an effective therapeutic strategy to enhance anti-tumor immune responses.

## Materials and Methods

### Mice studies

All the animal studies were conducted in compliance with the guidelines of the Institutional Animal Care and Use Committee (IACUC) at the University of Michigan. Arg1^f/f^ mice (Stock # 008817) and C57BL/6 wild-type (WT) mice (Stock# 000664) were obtained from the Jackson Laboratory and bred in-house. Arg1^f/f^ mice were generated to have *loxP* sites flanking exons 7 and 8 in the *Arg1* gene(El Kasmi et al., 2008). LysM^Cre/+^ mice and KF (Ptf1a^FlpO/+^;Kras^Frt-STOP-Frt-G12D/+^) mice were donated by Dr. Howard Crawford. LysM^Cre/+^ mice express Cre in myeloid cells due to the insertion of the Cre cDNA into the endogenous M lysozyme (LysM) locus^4^. LysM^Cre/+^;Arg1^f/f^ mice were generated by crossing LysM^Cre/+^ mice with Arg1^f/f^ mice. KFCA mice were generated by crossing KF mice with LysM^Cre/+^;Arg1^f/f^ mice. For the transplantation mouse model, we injected 50,000 7940B cells (C57BL/6J) into C57BL/6J mouse pancreas. 7940B cells were derived from a male KPC (P48-Cre; loxP-stop-loxP (LSL)-Kras^G12D^; p53^flox/+^) mouse tumor(Long et al., 2016).

### Chemical compounds

Arginase inhibitor, CB-1158 was synthesized and provided by Calithera Biosciences, Inc., South San Francisco, CA (2017). For the mice studies, CB-1158 was dissolved in Milli-Q water and administered by oral gavage twice a day at 100 mg/kg. This treatment started 10 days after tumor implantation and lasted for 10 days. The control group received Milli-Q water by oral gavage, twice a day. Purified anti-mouse PD1 antibody (BioXcell #BE0033-2; clone J43) was used for the *in-vivo* anti-PD1 blockade experiments. Anti-PD1 was used at a dose of 200 μg/i.p. injection, every 3 days. The control group received Polyclonal Armenian hamster IgG (BioXcell, BE0091) and it was administered in parallel to anti-PD1.

### Single-cell RNA-seq

Human sc-RNA-seq data were previously published in (N. G. Steele et al., 2020) (NIH dbGaP database accession #phs002071.v1.p1). Healthy mouse pancreas sc-RNA-seq data were previously published in (Kemp, Steele, et al., 2021) (NIH dbGap database accession #GSM5011581) and mouse spontaneous PDA sc-RNA-seq data generated using the KPC (*Kras^LSL-G12D^; Trp53^LSL-R172H^; Ptf1a-Cre*) model were previously published (NIH dbGap databse accession GSE202651). To generate the KF and KFCA sc-RNA-seq data, pancreatic tissue was harvested from KF (n=1) and KFCA mice (n=1) at 11 months of age. The tissue was mechanically minced, then digested with Collagenase V (Sigma C9263, 1mg/ml in RPMI) for 30 minutes at 37°C with shaking. Digestions were filtered through 500 μm, 100 μm, and 40 μm mesh to obtain single cells. Dead cells were removed using the MACS Dead Cell Removal Kit (Miltenyi Biotec). Single-cell complementary DNA libraries were prepared and sequenced at the University of Michigan Advanced Genomics Core using the 10x Genomics Platform. Samples were run using 50-cycle paired-end reads on the NovaSeq 6000 (Illumina) to a depth of 100,000 reads. The raw data were processed and aligned by the University of Michigan Advanced Genomics Core. Cell Ranger count version 4.0.0 was used with default settings, with an initial expected cell count of 10,000. Downstream sc-RNA-seq analysis was performed using R version 4.0.3, R package Seurat version 4.0.2, and R package SeuratObject version 4.0.1 (RStudio Team RStudio: Integrated Development for R (RStudio, 2015); http://www.rstudio.com/ R Core Development Team R: A Language and Environment for Statistical Computing (R Foundation for Statistical Computing, 2017); https://www.R-project.org/ (Butler, Hoffman, Smibert, Papalexi, & Satija, 2018; Stuart et al., 2019).Data were filtered to only include cells with at least 100 genes and genes that appeared in more than three cells. Data were normalized using the NormalizeData function with a scale factor of 10,000 and the LogNormalize normalization method. Data were then manually filtered to exclude cells with <1000 or >60,000 transcripts and <15% mitochondrial genes. Variable genes were identified using the FindVariableFeatures function. Data were scaled and centered using linear regression of transcript counts. PCA was run with the RunPCA function using the previously defined variable genes. Cell clusters were identified via the FindNeighbors and FindClusters functions, using dimensions corresponding to approximately 90% variance as defined by PCA. UMAP clustering algorithms were performed with RunUMAP. Clusters were defined by user-defined criteria. The complete R script including figure-specific visualization methods is publicly available on GitHub (https://github.com/PascaDiMagliano-Lab/).

### Kaplan-Meier survival analysis

For Kaplan-Meier overall survival, we used the human dataset GSE71729 containing 125 primary PDA tumor samples. The samples were split into *ARG1*-low (n=62) and *ARG1*-high (n=63) groups. Survival analysis with log-ranked test was subsequently plotted in GraphPad Prism v9.

### Histopathology

Tissues were fixed in 10% neutral-buffered formalin overnight, dehydrated, paraffin-embedded and sectioned into slides. Hematoxylin and Eosin (H&E) and Gomori’s Trichrome staining were performed according to the manufacturer’s guidelines.

#### Immunohistochemistry (IHC)

Paraffin sections were re-hydrated with 2 series of xylene, 2 series of 100% ethanol and 2 series of 95% ethanol. Slides were rinsed with water to remove previous residues. CITRA Plus (BioGenex) was used for antigen retrieval and microwaved for 5 minutes and then 3 minutes. Once cool down, sections were blocked with 1% bovine serum albumin (BSA) in PBS for 30 minutes. Primary antibodies were used at their corresponding dilutions (Supplementary Table) and incubated at 4°C overnight. Biotinylated secondary antibodies were used at a 1:300 dilution and applied to sections for 45 minutes at room temperature. Sections were then incubated for 30 minutes with ABC reagent from Vectastain Elite ABC Kit (Peroxidase), followed by DAB (Vector).

#### Co-immunofluorescence (Co-IF)

Deparaffinized slides were blocked with 1% BSA in PBS for 1 hour at RT. Primary antibodies (Supplementary Table) were diluted in blocking buffer and incubated overnight at 4°C, followed by secondary antibody (Alexa Fluor secondaries, 1:300) for 45 minutes at RT. Slides were mounted with Prolong Diamond Antifade Mountant with DAPI (Invitrogen). TSA Plus Fluorescein (PerkinElmer) was also used in the Co-IF for primary antibodies.

#### in situ hybridization (ISH) with Co-IF

The RNA Scope Multiplex Fluorescent Detection Kit (Advanced Cell Diagnostics) was used according to the manufacturer’s protocol. The probe used for *ARG1* was Hs-ARG1(401581, Advanced Cell Diagnostics). Freshly cut human paraffin-embedded sections were baked for 1 hour at 60°C prior to staining. Slides were then deparaffinized and treated with hydrogen peroxide for 10 minutes at room temperature. Target retrieval was performed in a water steamer boiling for 15 minutes, and then slides were treated with the ProteasePlus Reagent (Advanced Cell Diagnostics) for 30 minutes. The RNA scope probe was hybridized for 2 hours at 40°C. The signal was amplified using the AMP materials provided in the ACD Multiplex Kit (Advanced Cell Diagnostics). The signal was developed with horseradish peroxidase (HRP) channel. Once completed, the samples were washed in PBS, and then blocked for 1 hour with 5% donkey serum at room temperature. Primary antibody against CD45 (1:400) was incubated overnight at 4°C. Secondary antibodies (1:300 in blocking buffer) were incubated for 1 hour at room temperature, and samples were washed three times in PBS. Slides were counterstained with DAPI and mounted with ProLong Gold Antifade Mountant (Thermo Fisher Scientific).

Images were taken either with an Olympus BX53 microscope, a Leica SP5 microscope, a Leica STELLARIS 8 FALCON Confocal Microscopy System, or scanned with a Pannoramic SCAN scanner (Perkin Elmer). Quantification of positive cell number or area was done using ImageJ, 3-5 images/slide (200x or 400x magnification) taken from 3-4 samples per group or using the Halo software (Indica Labs).

### Cell culture

The 7940B cells were cultured in Dulbecco’s Modified Eagle Medium (DMEM, 11965-092) supplemented with 10% Fetal Bovine Serum and 1% Penicillin Streptomycin. Tumor cell conditioned media (CM) were collected from the 7940B cells that were cultured to confluency. Media were centrifuged at 300 *g* for 10 minutes at 4°C to remove contaminating tumor cells. These CM were used for the macrophage polarization assay. For macrophage polarization, bone marrow (BM) cells were isolated from WT or LysM^Cre/+^;Arg1^f/f^ mice femurs. Once isolated, BM cells were cultured with 7940B cell CM plus total DMEM media at a 1:1 ratio for 6 days. Fresh media was added during day 3. This process allowed the differentiation and polarization of BM cells to TAMs(Zhang, Velez-Delgado, et al., 2017).

### Western Blot

TAMs from WT or LysM^Cre/+^;Arg1^f/f^ mice were lysed in RIPA buffer (Sigma-Aldrich) with protease and phosphatase inhibitors (Sigma-Aldrich). Protein samples were quantified, normalized, and then electrophoresed in a 4-15% SDS-PAGE gel (BioRad). Protein was transferred to a PVDF membrane (BioRad), blocked with 5% milk for one hour at room temperature, and then incubated with primary antibodies overnight at 4°C (Supplementary Table). Membranes were then incubated with HRP-conjugated secondary antibody (1:5000) for 2 hours at room temperature. Membranes were washed, incubated in Western Lightning Plus-ECL (PerkinElmer), and then visualized with the ChemiDoc Imaging System (BioRad).

### Metabolomics analysis

Conditioned media (200 µL) was collected from each well of WT and LysM^Cre/+^;Arg1^f/f^ TAMs in a 6-well plate after 6 days of culture and used for extracellular metabolite profiling. Briefly, to the 200 µL of media, 800 µL of ice-cold 100% methanol was added. Cell lysates from parallel plates were used for protein quantification, and the protein amount was used to normalize the volume of samples collected for metabolomics. The samples were centrifuged at 12,000 *g* for 10 minutes after which the supernatant was collected, dried using SpeedVac Concentrator, reconstituted with 50% v/v methanol in water, and analyzed by targeted liquid chromatography tandem mass spectrometry (LC-MS/MS) and processed as previously described(Nelson et al., 2020).

### Statistics

GraphPad Prism 9 was used to performed statistical analysis. T-test or ANOVA was performed for group comparisons. R software 3.5.2 was used for the analysis of the microarray data set. A p-value was considered statistically significant when p<0.05.

### Data Availability

Human sc-RNA-seq data was previously published(N. G. Steele et al., 2020) and both raw and processed data are available at the NIH dbGap database accession number phs002071.v1.p1. Raw and processed sc-RNA-seq data for the WT and KPC were previously published and are available at GEO accession number GSM5011580 and GSE202651. Raw and processed sc-RNA-seq data for the KF and KFCA are available at GEO accession number GSE203016.

## Acknowledgements

We thank Daniel Long and Michael Mattea for histology services at the University of Michigan. We thank Calithera Biosciences for providing the Arginase inhibitor, CB-1158. We thank the Advanced Genomics core at the University of Michigan for RNA sequencing. We also thank the Microscopy Core and the Flow Cytometry Core at the University of Michigan Biomedical Research Core Facilities for providing access to advanced microscopy and flow cytometry. We thank the Microscopy, Imaging and Cellular Physiology Core of the Michigan Diabetes Research Center at the University of Michigan for providing access to the Leica STELLARIS 8 FALCON Confocal Microscopy System, funded by grant NIH S10OD28612-01-A1.

## Notes

**Funding:** This study was supported by NIH U01CA224145, NIH/NCI R01CA151588, R01CA198074, and by an American Cancer Society Scholar grant to M. Pasca di Magliano. This study was also supported by the University of Michigan Cancer Center Support Grant (NCI P30CA046592), including an Administrative Supplement to M. Pasca di Magliano. C.A. Lyssiotis was supported by the NCI (R37CA237421, R01CA248160, R01CA244931). H.C. Crawford was supported by R01 CA247516. R.E. Menjivar was supported by the University of Michigan Rackham Merit Fellowship, by the Cellular and Molecular Biology Training Grant (NIH T32-GM007315), by the Center for Organogenesis Training Program (NIH T32-HD007505), and by the NCI (F31-CA257533). Z.C. Nwosu was supported by the Michigan Postdoctoral Pioneer Program, University of Michigan Medical School. WD was supported by University of Michigan Training Program in Organogenesis. K.L. Donahue was supported by the Cancer Biology Program training grant (T32-CA009676). A. Velez-Delgado was supported by the Rackham Merit Fellowship, by the Cellular Biotechnology Training Program (T32GM008353) and by the NCI (F31-CA247037). P. Kadiyala was supported by the Immunology training grant (T32-AI007413). D. Salas-Escabillas was supported by the Rackham Merit Fellowship and by the Cancer Biology Program training grant (T32-CA009676). E.S. Carpenter was supported by the American College of Gastroenterology Clinical Research Award and by T32-DK094775. Y. Zhang was funded by the NCI-R50CA232985. C.J Halbrook was supported by F32CA228328, K99/R00CA241357, P30CA062203, and the Sky Foundation. The funders did not have a role in the planning, execution, or writing of this study.

**Competing Interests:** C.A.L. has received consulting fees from Astellas Pharmaceuticals, Odyssey Therapeutics, and T-Knife Therapeutics, and is an inventor on patents pertaining to Kras regulated metabolic pathways, redox control pathways in pancreatic cancer, and targeting the GOT1-pathway as a therapeutic approach (US Patent No: 2015126580-A1, 05/07/2015; US Patent No: 20190136238, 05/09/2019; International Patent No: WO2013177426-A2, 04/23/2015).

### Competing Interest Statement

C.A.L. has received consulting fees from Astellas Pharmaceuticals, Odyssey Therapeutics, and T-Knife Therapeutics, and is an inventor on patents pertaining to Kras regulated metabolic pathways, redox control pathways in pancreatic cancer, and targeting the GOT1-pathway as a therapeutic approach (US Patent No: 2015126580-A1, 05/07/2015; US Patent No: 20190136238, 05/09/2019; International Patent No: WO2013177426-A2, 04/23/2015).

