## Supplemental Figures for "Arginase 1 is a key driver of immune suppression in pancreatic cancer"

### Figure 1- figure supplement 1

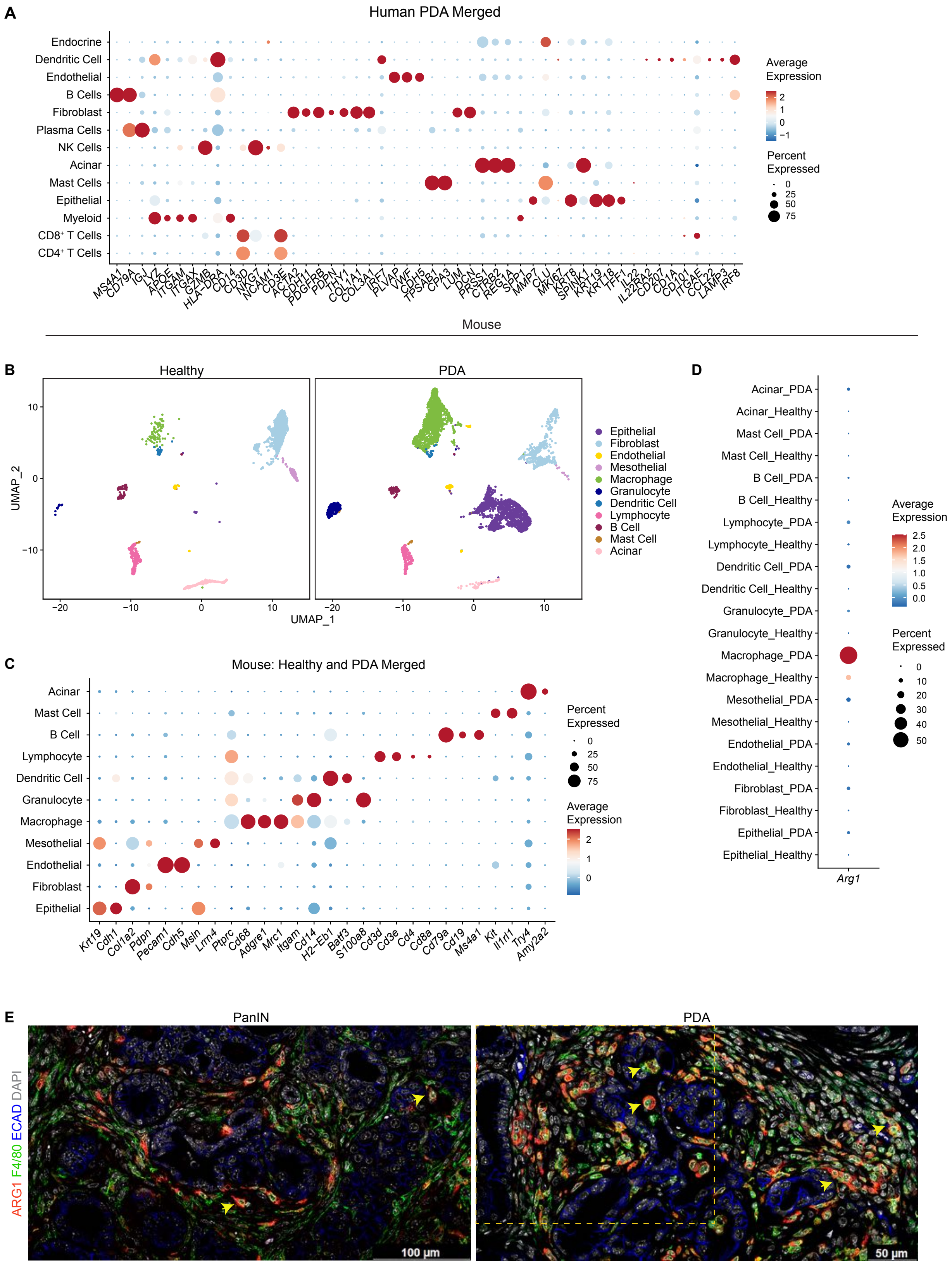

**Arg1 expression in human and mouse myeloid cells.** (**A**) Dot plot showing the lineage markers used to identify the different cell populations in the human Sc-RNA-seq. Dot color indicates average expression and dot size indicates expression frequency. (**B**) UMAP visualizations comparing defined cell populations in healthy and PDA mouse sc-RNA-seq. (**C**) Dot plot showing the lineage markers used to identify the distinct cell populations in healthy and PDA mouse sc-RNA-seq. (**D**) Dot plot showing normalized gene expression levels of *Arg1* in healthy and PDA mouse sc-RNA-seq. (**E**) Representative co-immunofluorescence staining for ARG1 (red), macrophages (F4/80, green), and epithelial (ECAD, blue) cells in mouse tissue during PanIN and PDA stages of disease. Yellow dotted box marks the area shown in main Figure 1H. Scale bar for PanIN image is 100 μm; for PDA is 50 μm.

### Figure 2- figure supplement 1

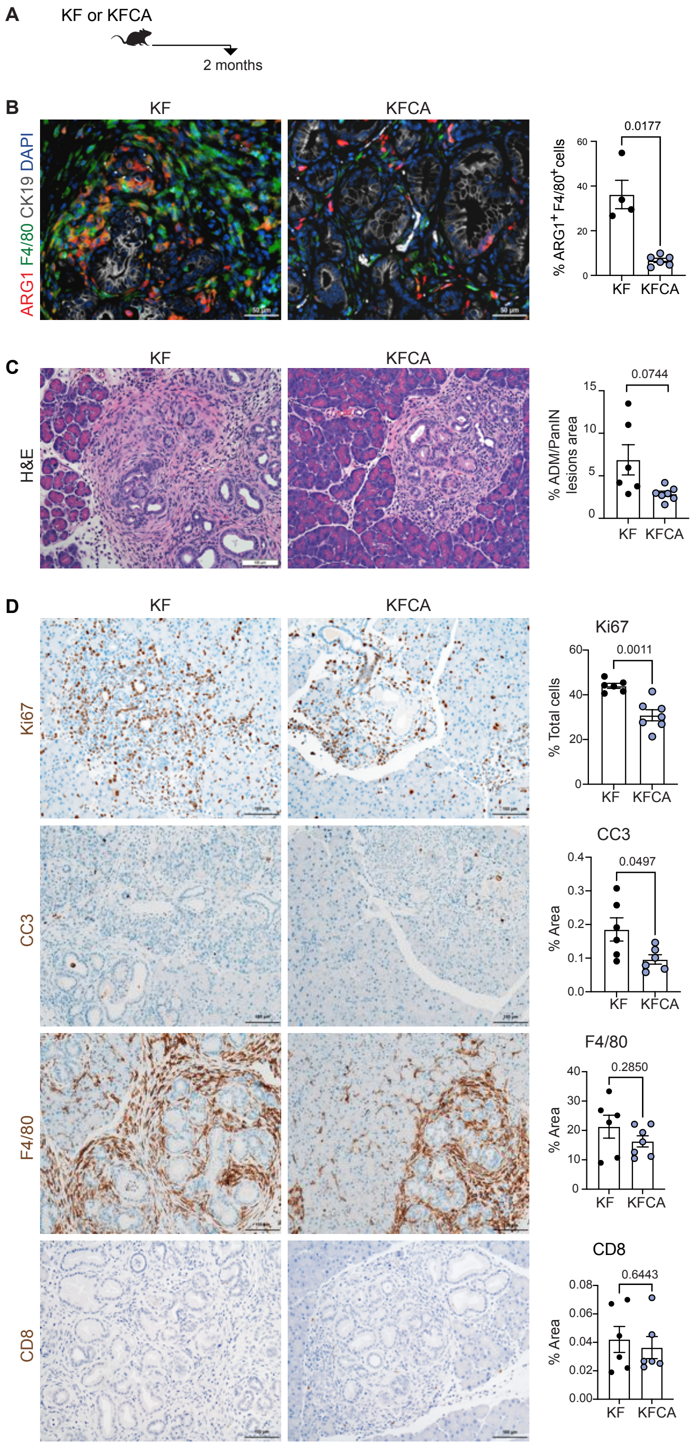

**Deletion of Arg1 in myeloid cells decreases cell proliferation and cell death during early stages of PDA.** (**A**) Experimental design for KF and KFCA mice aged to 2 months. (**B**) Representative images of co-immunofluorescence staining for ARG1 (red), macrophages (F4/80, green), epithelial (CK19, gray), and DAPI (blue) in KF and KFCA pancreata at 2 months. Yellow staining shows co-localization of ARG1 and F4/80. Scale bar, 50 μm. Quantification on the right, n= 4-6/group (**C**) Representative H&E images for KF and KFCA mice at 2 months, n=6-7/group. Scale bar, 100 μm. On the right, percentage of ADM/PanIN lesions area in KF and KFCA tissue at 2 months, n=6-7/group. (**D**) Representative immunohistochemistry staining for cell proliferation (Ki67), cell death (CC3), macrophages (F4/80), and $CD8^+$ T cells (CD8) in KF and KFCA tissue at 2 months. Scale bar, 100 μm. Positive staining quantification is shown on the right. Student's t test was used to determine significance.

### Figure 2- figure supplement 2

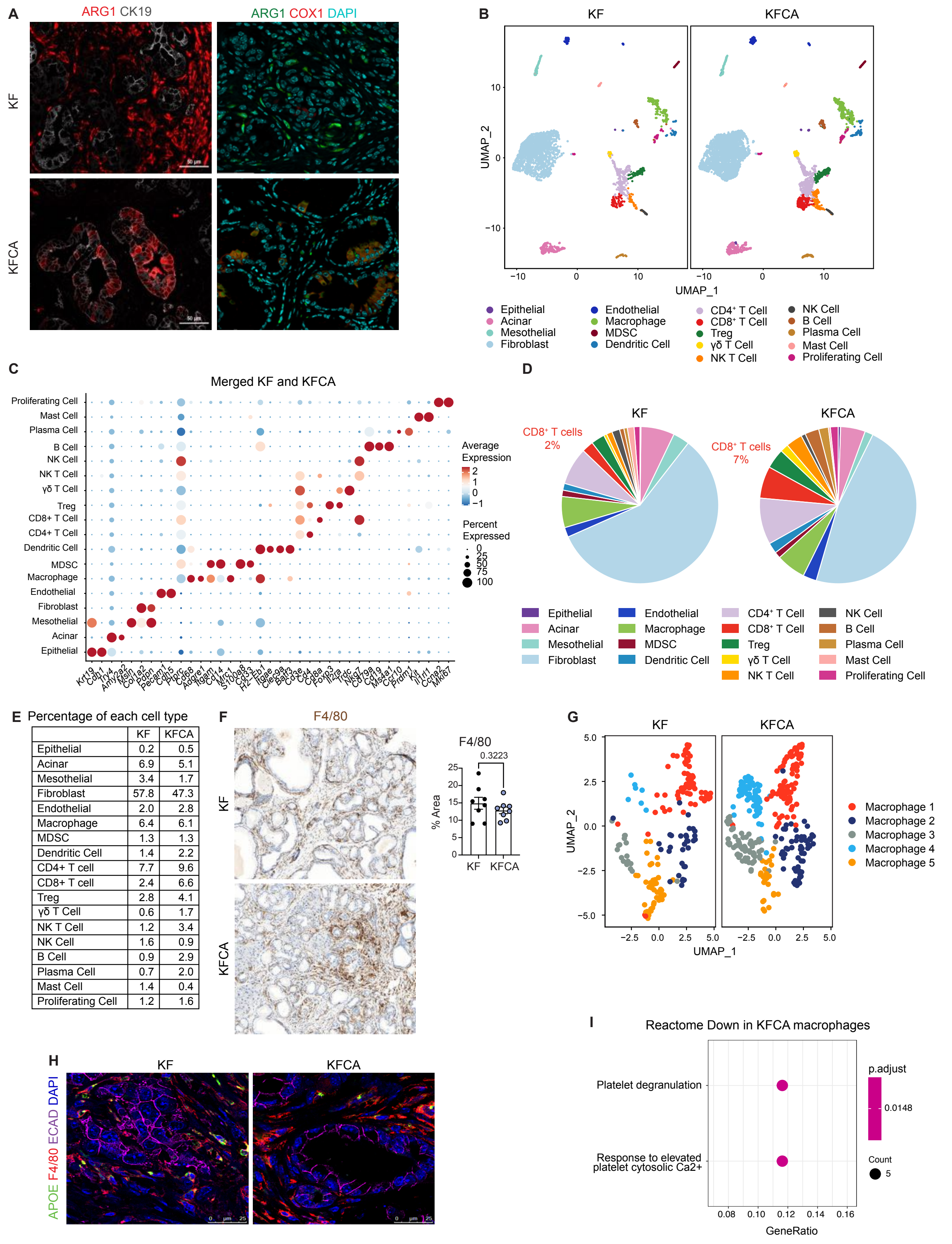


| | KF | KFCA |
|---|---|---|
| Epithelial | 0.2 | 0.5 |
| Acinar | 6.9 | 5.1 |
| Mesothelial | 3.4 | 1.7 |
| Fibroblast | 57.8 | 47.3 |
| Endothelial | 2.0 | 2.8 |
| Macrophage | 6.4 | 6.1 |
| MDSC | 1.3 | 1.3 |
| Dendritic Cell | 1.4 | 2.2 |
| CD4+ T cell | 7.7 | 9.6 |
| CD8+ T cell | 2.4 | 6.6 |
| Treg | 2.8 | 4.1 |
| γδ T Cell | 0.6 | 1.7 |
| NK T Cell | 1.2 | 3.4 |
| NK Cell | 1.6 | 0.9 |
| B Cell | 0.9 | 2.9 |
| Plasma Cell | 0.7 | 2.0 |
| Mast Cell | 1.4 | 0.4 |
| Proliferating Cell | 1.2 | 1.6 |

**Arginase 1 deletion in myeloid cells induces immune suppression in a spontaneous PDA mouse model during late stages of disease.** (**A**) Left panel: Co-immunofluorescence staining from Figure 2F showing ARG1 (Red) and CK19 (gray) staining without DAPI; right panel: co-immunofluorescence staining for ARG1 (green), COX1 (Red) and DAPI (Teal) in KF and KFCA tissue. Scale bar, 50 µm. (**B**) UMAP visualizations of sc-RNA-seq comparing the identified cell populations in KF and KFCA. (**C**) Dot plot showing the lineage markers used to identify the distinct cell populations. Dot color indicates average expression and dot size indicates expression frequency. (**D**) Pie charts comparing the proportion of the identified sc-RNA-seq populations between KF and KFCA. (**E**) Table showing the percentage of each identified population from total cells in KF and KFCA. (**F**) Representative immunohistochemistry staining for macrophages (F4/80) in KF and KFCA tissue during late stage of disease at 20x magnification, n=8/group. Positive staining quantification is shown to the right. Student's t test was used to determine significance. (**G**) UMAP visualizations showing the distribution of the 5 identified macrophage subpopulations in KF and KFCA. (**H**) Representative immunofluorescence staining for APOE (green), F4/80 (red), ECAD (Purple) and DAPI (blue) in KF and KFCA pancreas tissue during late stage of disease. Scale bar, 25 µm. (**I**) Reactome pathway enrichment analysis showing significantly downregulated pathways in KFCA macrophages.

#### Figure 4- figure supplement 1

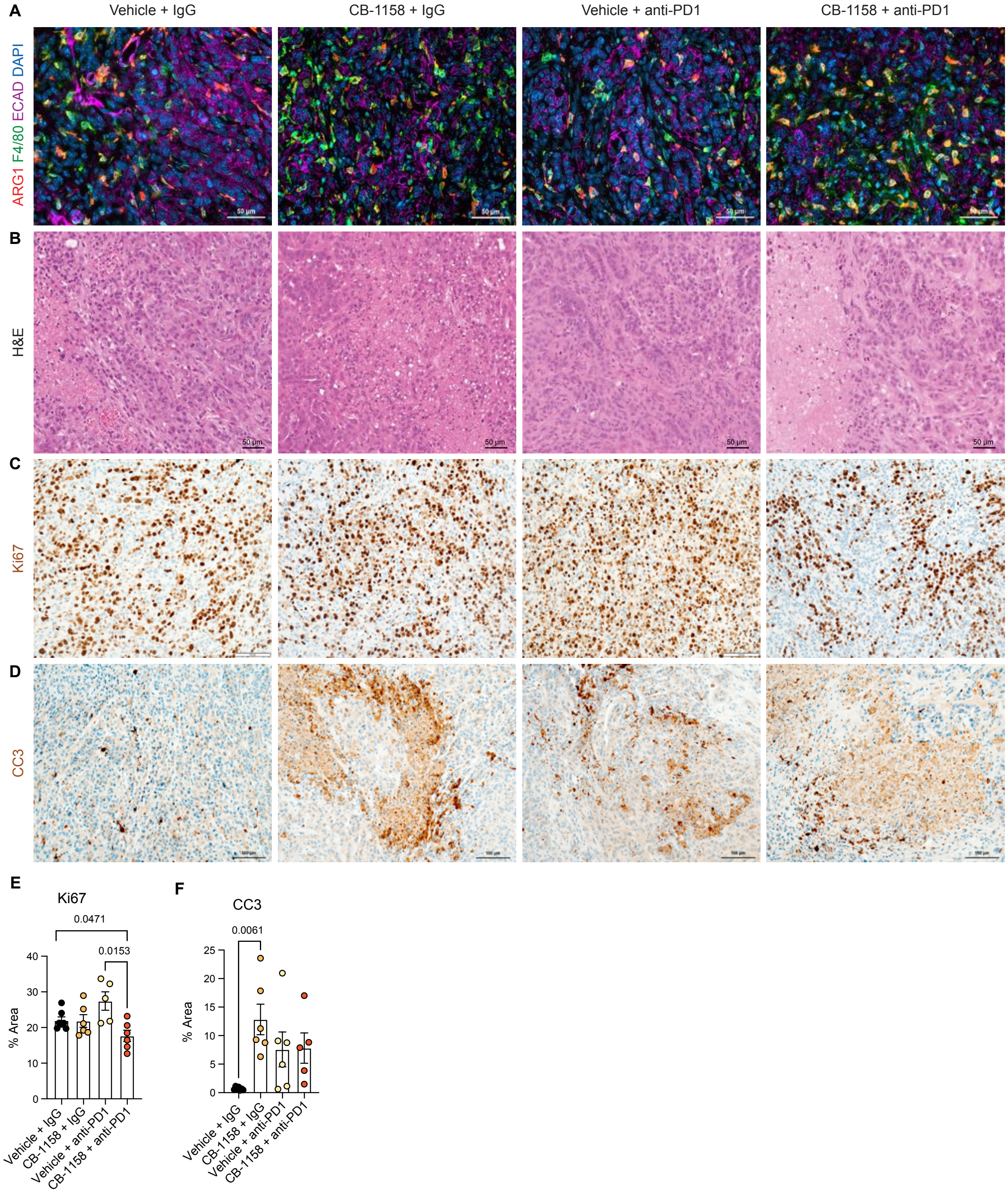

**Histological analysis of the multiple treatment groups in the orthotopic mouse model.** (**A**) Representative images of co-immunofluorescence staining for ARG1 (red), F4/80 (green), ECAD (purple), and DAPI (blue). Yellow shows co-localization of ARG1 with F4/80 macrophages. Scale bar, 50 μm. (**B**) Hematoxylin and Eosin (H&E) staining for the indicated treatment groups. Scale bar, 50 μm. (**C**) Immunohistochemistry staining for cell proliferation (Ki67). Positive staining is shown in brown. Scale bar, 100 μm. (**D**) Immunohistochemistry staining for cell death (CC3). Positive staining is shown in brown. Scale bar, 100 μm. (**E** and **F**)**.** Quantification of the positive staining area for Ki67 and CC3 from **C** and **D.** Significance was determined using unpaired t test with Welch's correction.
