## Supplemental Table for "Arginase 1 is a key driver of immune suppression in pancreatic cancer"

**Supplementary Table. Immunohistochemistry, immunofluorescence, and western blot antibodies and dilutions.**

| Antibody | Supplier | Catalog Number | IHC dilution | IF dilution | WB dilution |
|---|---|---|---|---|---|
| ARG1 | Cell signaling | 93668S | | 1:75 | 1:1000 |
| CD45 | R&D Systems, Minneapolis, MN | MAB14302 | | 1:400 | |
| ECAD | Cell Signaling, Danvers, MA | 14472S | | 1:50 | |
| F4/80 | Cell signaling | 70076S | 1:250 | 1:250 | |
| CK19 (Troma III) | Developmental Studies Hybridoma Bank, Iowa City, IA | - | | 1:50 | |
| Ki67 | Abcam, Cambridge, UK | ab15580 | | 1:100 | |
| CC3 | Cell Signaling | 9661L | 1:300 | | |
| CD8 | Cell Signaling | 98941S | 1:300 | 1:400 | |
| APOE | Abcam | ab183597 | | 1:500 | |
| COX1 | Santa Cruz | Sc-1754 | | 1:200 | |
| GZMB | Cell signaling | | | 1:800 | |
| Vinculin | Cell signaling | 13901S | | | 1:1000 |
